# Unexpectedly high coral heat tolerance at thermal refugia

**DOI:** 10.1101/2023.06.16.545328

**Authors:** Liam Lachs, Adriana Humanes, Peter J Mumby, Simon D. Donner, John Bythell, Elizabeth Beauchamp, Leah Bukurou, Daisy Buzzoni, Ruben de la Torre Cerro, Holly K. East, Alasdair J. Edwards, Yimnang Golbuu, Helios M. Martinez, Eveline van der Steeg, Alex Ward, James R. Guest

## Abstract

Marine heatwaves and mass bleaching have led to global declines in coral reefs. Corals can adapt, yet, to what extent local variations in thermal stress regimes influence heat tolerance and adaptive potential remains uncertain. Here we identify persistent local-scale thermal refugia and hotspots among the reefs of a remote Pacific archipelago, based on 36 years of satellite-sensed temperatures. Theory suggests that hotspots should promote coral heat tolerance through acclimatisation and directional selection. While historic patterns of mass bleaching and marine heatwaves align with this expectation, we find a contrasting pattern for a single species, *Acropora digitifera*, exposed to a marine heatwave experiment. Higher heat tolerance at thermal refugia (+0.7 °C-weeks) and correlations with other traits suggest that non-thermal selective pressures may also influence heat tolerance. We also uncover widespread heat tolerance variability, indicating climate adaptation potential. Compared to the least-tolerant 10% of the *A. digitifera* population, the most-tolerant 10% could withstand an additional heat stress of 5.2 and 4.1 °C-weeks for thermal refugia and hotspots, respectively. Despite expectations, local-scale thermal refugia can harbour higher heat tolerance, and mass bleaching patterns do not necessarily predict species responses. This has important implications for designing climate-smart initiatives to tackle global-scale adaptive management problems.

## 1. Introduction

Climatic stressors are causing profound impacts on organisms and ecological systems globally^1,2^. This is especially true for sessile organisms which cannot escape their environment. To persist under climate change, many species will need to adapt, and this will require sufficient standing genetic variability of climate resilience traits within populations^3^.

Over large geographic scales, gradients in climatic exposure are known to shape organism tolerance to climatic stressors^4,5^. However, whether these trends hold over the smaller spatial scales at which organisms and populations operate remains poorly understood, despite the crucial need for local-scale information for effectively managing populations under climate change.

Reef-building corals are the poster child for this problem because of their acute sensitivity to marine heatwave stress, which disrupts their symbiosis with phototrophic microalgae and can lead to mass bleaching and mortality^6–8^. Some coral reefs are routinely exposed to heat stress while others consistently avoid it, herein referred to as hotspots and thermal refugia, respectively. These distinct thermal stress regimes have been identified across broad geographic regions (>1000 km, *e*.*g*., the Great Barrier Reef^9^, Red Sea^4^, and Caribbean^10^). As for many other marine and terrestrial species, the thermal limit of corals largely tracks broad climatological gradients^7,11,12^. When exposed to marine heatwaves, corals at hotspots have tended to fair better than those at thermal refugia, through means of acclimatisation^13,14^ and adaptation^15,16^. From a management perspective, it is desirable if heterogeneity in thermal environments leads to predictable differences in coral population responses across seascapes, as this could allow managers to achieve higher efficacy through prioritising interventions in locations where they can be most effective. However, there is growing recognition that heat tolerance is highly variable within single populations^17^, which could influence the feasibility of spatial seascape management plans. Because management typically takes place over local spatial scales, there is a growing need to study smaller environmental gradients to test a critical unresolved question: does the large-scale theory that hotspots harbour higher levels of heat tolerance hold true over smaller manageable distances?

Here, we combine the concept that thermal gradients lead to spatial heterogeneity in coral heat tolerance, with a more detailed exploration of within-population variability of this trait. We ask whether the difference in heat tolerance between hotspots and thermal refugia observed over broad spatial scales is robust at the intrapopulation scale, given high levels of phenotypic variability. One reason that this large-scale concept may not apply to single populations is if tolerance to marine heatwaves is not only a function of the thermal environment in which corals live, but also influenced by other environmental drivers (*e*.*g*., water quality^18,19^) or biological factors (*e*.*g*., energy reserves^20^). Our study takes a local-scale (< 100 km) seascape perspective using historic patterns of mass coral bleaching, and a 5-week simulated marine heatwave experiment on a common coral species from multiple reefs at hotspots and thermal refugia. If we find that the broad-scale theory is robust within populations, then it simplifies the provision of advice to ecosystem managers, that hotspot reefs might harbour corals with predictably higher levels of heat tolerance than thermal refugia.

## 2. Methods

To test whether persistent hotspots or thermal refugia are present in Palau, spatial patterns in thermal history and marine heatwaves were assessed using DHW. Firstly, the spatial variability of bleaching risk across the entire coral assemblage was tested using a Bayesian statistical modelling approach linking DHW data with historic bleaching survey observations from across Palau. Then we restricted our spatial comparison to a single species avoid any possible confounding effects of species compositional changes on coral responses to heat stress. We achieved this by conducting a long-term (5-week) marine heatwave experiment with assays of bleaching and mortality every 1–3 days.

### 2.2. Historic heat stress data

Heat stress on Palauan coral reefs was calculated from CoralTemp version 3.1, a 0.05 ° x 0.05 ° latitude-longitude resolution satellite-based Sea Surface Temperature (*SST*) dataset (1985 to 2020) available from the National Oceanic and Atmospheric Administration’s Coral Reef Watch (NOAA CRW)^51^. Although coral bleaching and mortality can be caused by light intensity, cold spells, nutrient enrichment, and sedimentation, among other factors, chronic levels of accumulated heat stress are the single factor most strongly associated with mass coral bleaching and mortality^6^. NOAA CRW measures accumulated heat stress with the Degree Heating Weeks metric (DHW) based on CoralTemp v3.1 (1985 to 2020). Coral reefs exposed to higher levels of DHW have a greater risk of mass bleaching and mortality^52^. As such, DHW is used by NOAA CRW to provide real-time bleaching risk forecasts, whereby DHW of 4–8 °C-weeks corresponds to the expectation of significant bleaching, and DHW > 8 °C-weeks corresponds to the expectation of significant bleaching and mortality. Here we follow the NOAA CRW methodology^21^, DHW on a given day (*i*) is computed as the sum of the last 12 weeks (84 days) of daily temperature anomalies (*HotSpots*) relative to a standard coral stress threshold baseline (*MMM*, maximum of monthly means) which is held constant through time. Only Hotspots > 1 °C are accumulated and are divided by 7 to make DHW a weekly metric.

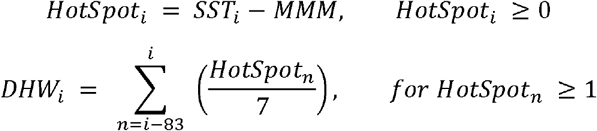

### 2.3. Identifying persistent hotspots and thermal refugia

Whether or not hotspots and thermal refugia were present in Palau was determined using timeseries of maximum annual heat stress statistics (DHW derived from CoralTemp), combined with a thermal consistency biplot analysis (Fig. 1). For each year independently, maps of DHW were transformed to percentile ranks to show the relative spatial differences in heat stress for all grid cells that intersect coral reefs in Palau^53,54^. To describe the relative thermal history of each reef, we calculated their typical heat stress ranking (average rank) and rank consistency (rank SD) through time. As some reefs are always more or less heat stressed in comparison to others, the most- and least-heat stressed reefs in Palau could be easily determined using typical heat stress ranks based on the upper and lower terciles.

**Figure 1.**
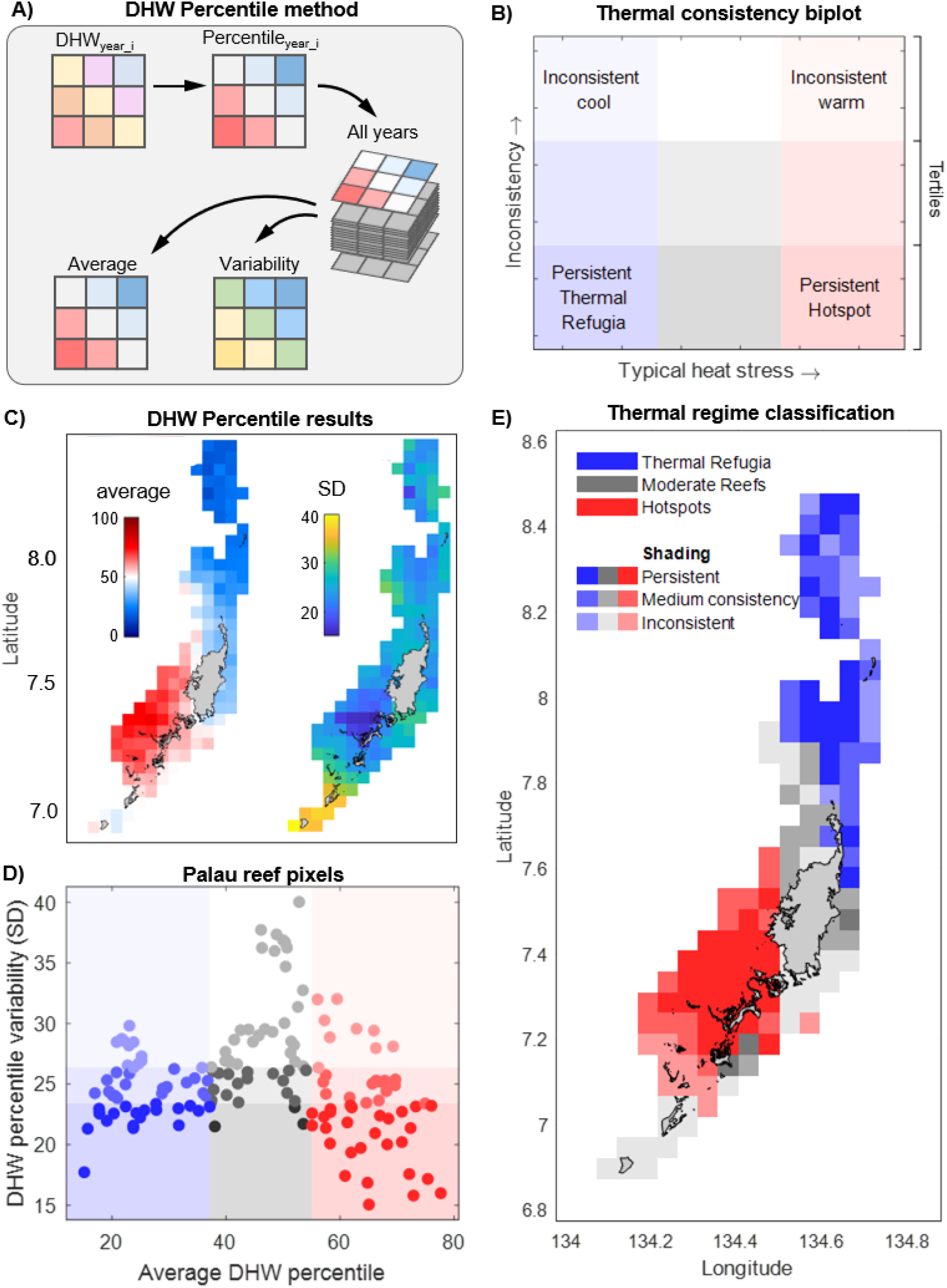
Characterisation of thermal regimes across Palau showing conceptual diagrams (A-B) and empirical results (C-E). (A) Time series of heat stress percentiles based on Degree Heating Weeks (DHW) are used to determine spatial patterns in (C) the typical heat stress (average DHW percentile) and heat stress consistency (DHW percentile variability in standard deviations – SD). The thermal consistency biplot (C – theory, D – Palau) defines coral reefs according to these two metrics, where equal-sized tercile subsets of each variable demarcate distinct thermal regimes. (E) Persistent thermal refugia are located among the northern reefs, while persistent hotspots dominate in the southwest.

However, to be considered hotspots or thermal refugia, reefs must have such thermal conditions consistently through time. Therefore, the reefs were also categorised based on DHW percentile SD, with the most-consistent thermal regimes occurring in the lower tercile, and the least-consistent thermal regimes occurring in the upper tercile. Using the thermal consistency biplot analysis, which plots typical heat stress rank against consistency, the tercile groups resulted in 9 regions. The lower left and lower right terciles of the biplot (Fig. 1B, 1C) thus indicate the most-persistent thermal refugia and hotspots in the study region. The historic DHW trends for these regions were further inspected, confirming that bleaching-level heat stress conditions occurred far more often in the hotspots, while non-heat stress years were far more frequent in the thermal refugia.

### 2.4. Assemblage-wide coral bleaching data

Coral bleaching is caused by a breakdown in the symbiosis between coral hosts and their photosynthetic algal symbionts and often leads to coral mortality. Underwater coral bleaching survey data used in this study were collected between 1998 and 2017, encompassing the two major bleaching events in Palau (1998 and 2010). This dataset is publicly available^55,56^ and provides the timing and coordinates of the survey. To account for multiple survey methods in the data collection, bleaching observations (primarily reported as percentage of reef area with bleached corals) were summarised as severity scores ranging from 0 to 3, namely: no bleaching (0%), mild bleaching (1-10%), moderate bleaching (11-50%), and severe bleaching (>50%).

### 2.5. Spatial variability in assemblage-wide coral bleaching

To quantify the spatial variability of DHW-bleaching responses across the coral assemblage (*i*.*e*., all species) in Palau, we combined CoralTemp with historic coral bleaching survey observations. The effect of DHW on bleaching was fitted using a spatial beta GLM via Integrated Nested Laplace Approximation (INLA) in R-INLA. This Bayesian statistical approach provides a more intuitive spatially explicit estimation of model uncertainty^57,58^.

Spatial dependencies in bleaching observations are dealt with by implementing the Matérn correlation across a Gaussian Markov random field (GMRF). Essentially, this is a map of spatially correlated uncertainty that shows where bleaching is under-or over-predicted for a given heat stress dosage (DHW). The parameters (Ω) that determine the Matérn correlation are the range (*r* – range at which spatial correlation diminishes) and error (σ). To fit the beta distribution, bleaching severity scores were converted to proportions (divide by 3) and transformed to remove extremes of 0 and 1, which in practice is (Y x (n -1) + 0.5)/n, where *y* is the bleaching severity score scaled from 0 to 1, and *n* is the sample size^59^. This approach preserves differences among bleaching categories that would be lost using other transformations (*e*.*g*., binary transformation for a binomial GLM^22^).

Spatial variation in the uncertainty of bleaching-DHW responses was estimated across a high-resolution Delaunay triangulation mesh of the study area with a maximum triangle edge length of 2 km and a low-resolution convex hull (convex = -0.1) around the study sites to avoid boundary effects (6,210 nodes). The probability of severe coral bleaching for a given observation (*CB*_*i*_) and location (*i*) was assumed to follow a beta distribution (fitted values of π_*i*_, and precision parameter θ) using the logit-link function. Bleaching was modelled as a function of heat stress (*DHW*_*i*_) whilst accounting for additional underlying spatial correlation among bleaching observations (random effect: *u*_*i*_),

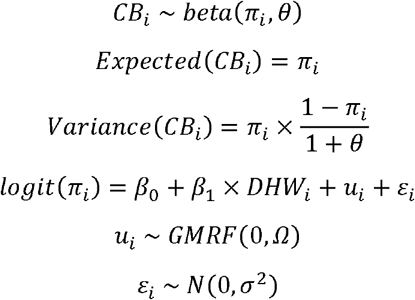

where β_*0*_ is the intercept, β_*1*_ is the slope for DHW, *u*_*i*_ represents the smoothed spatial effect from the GMRF mesh, elements of Ω (*r* and σ) are estimated from the Matérn correlation, and ε_*i*_ contains the independently distributed residuals. We specified weakly informative priors for *r*_*0*_ (20 km) and σ_*0*_ (1) based on the residual variogram and recommendations from ^58^, respectively. We also tested different priors; however, they had a negligible effect on the estimates of any model parameters. Whether DHW-based predictions of bleaching severity (u values) were significantly different between thermal refugia and hotspots was tested using a Wilcoxon sum rank test un the subset of nodes that intersected with the area of these thermal regimes as determined through the satellite data analysis described previously.

### 2.6. Simulated marine heatwave experiment

Natural selection and evolution occur within species and are likely to be strongly reliant on individual heat tolerance. To address this, we quantified how thermal regimes influence HT by conducting a long-term (*sensu* ^60^) 5-week simulated marine heatwave experiment, inducing bleaching and mortality in 1020 replicate fragments from 178 colonies of *Acropora digitifera*, a widely distributed reef-building coral. Fragments from 90 colonies were collected from 6 replicate reefs located in hotspot grid cells (3 sites) and refugia grid cells (3 sites), with at least 4 km between sites. In comparison to short heat-shock experiments that typically last 1-2 days, this experimental temperature profile (Table S1) was designed to match the intensity and duration of a natural marine heatwave more closely^17,60^, with the assumption that the phenotypic bleaching and mortality responses would be more ecologically relevant. After collection, fragments were acclimatised for 7–10 days in tanks maintained at ambient sea temperature. At the start of the experiment, the temperature in stress tanks was increased gradually over two weeks, reaching a final bleaching-level temperature of approximately + 2.5 ºC above the site-specific climatological stress accumulation thresholds (MMM_adj_ + 1 ºC). This temperature was maintained for the remainder of the experiment (Fig. S1, Table S1). The use of flow-through tank systems allowed an element of natural diel temperature variability^61^. Calibrated HOBO loggers were placed in each tank and recorded temperatures at 10-minute intervals. To relate coral bleaching and mortality responses to accumulated heat stress, not instantaneous temperature, we calculated heat stress in the experiment using the Degree Heating Weeks (DHW) metric. To compare our experimental DHWs for each site with the NOAA coral bleaching forecasts (satellite-derived DHW), we used local *in situ* temperature data to adjust the satellite-based CoralTemp baselines (MMM – maximum of monthly means climatology – from the 0.05 ° grid cell encompassing a collection site), following ^17^. This involved adjusting the MMM based on the linear regression between daily 0.05° sea surface temperature (CoralTemp v3.1) and daily averaged *in situ* temperatures (recorded from additional HOBO loggers deployed at each collection site). This produced six adjusted local climatological baselines (MMM_adj_ – adjusted MMM), one for each collection site.

### 2.7. Heat tolerance

Since corals can recover from bleaching, a pragmatic definition of heat tolerance is a coral’s ability to survive levels of heat stress sufficient to cause bleaching and mortality in nature^62,63^. Fragments (6 per colony) were dispersed in random locations among ten heat stress tanks (5 fragments/colony) and two procedural control tanks (1 fragments/colony), such that there was an even distribution of site and colony replication among all tanks. If a fragment died in a procedural control tank, held at non-stressful ambient temperature conditions, it was an indication of handling effects for that colony, so all remaining fragments from the colony were removed from the experiment and the colony was not assigned a heat tolerance score. The health status of each fragment was scored visually at intervals of between 1 and 3 days. We used the bleaching and mortality index (*BMI*)^17,64^ to categorise coral bleaching and mortality responses, such that c_1_ to c_5_ are the proportion of replicate fragments (per colony) recorded as healthy (*c*_1_), half bleached (*c*_2_), bleached (*c*_3_), partial mortality (*c*_4_), or dead (*c*_5_), and *N* is the total number of categories (here *N* = 5). For example, a colony whose replicate fragments are either all healthy or all dead at a certain time point, will have a BMI value of 0 or 1, respectively.

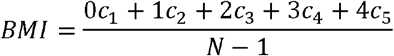

The BMI of a particular colony is representative of only a single timepoint and will change throughout the heat stress exposure. Therefore, we calculated DHW_50_, a colony-specific metric of heat tolerance analogous to the effective dosage (EC_50_) metric used in toxicology to determine the dosage at which 50% of the population experiences an effect. DHW_50_ for each colony was calculated by fitting a logistic dose response curve for all colonies, allowing a different intercept and slope per colony, and then computing the DHW value at which the fitted curve reaches 50% of the full bleaching and mortality response (*i*.*e*., a BMI value of 0.5) using the ‘qlogis’ function in R. This builds on coral heat stress experiments using the inverse average BMI through time as a measure of heat tolerance (1 – mean BMI). To allow for comparisons with previous work, we also computed the regression between inverse mean BMI and DHW_50_ (Fig. S2, *R*^*2*^ = 0.94).

### 2.8. Colony size and tissue biomass

For each *A. digitifera* colony, a downward-facing image was taken prior to sampling with a ruler for scale. These images were then analysed using the SizeExtractR workflow^65^ to determine the geometric mean diameter of each colony based on the maximum and minimum perpendicular diameter. Three additional fragments were sampled for each fragment to determine tissue biomass, as this is known to have an influence on energy reserves and potentially the ability to survive extended marine heatwave stress ^20^. Total tissue biomass for each fragment was determined as ash-free dry weight, or the difference between dry weight and ash weight. To achieve this, fragments were dried in an oven at the Palau International Coral Reef Center at 60 °C until constant weight (approximately 3 days). Branch tips were removed to approximate a cylinder, and the total length and midpoint width were recorded using vernier callipers. Each branch was crushed using a sterilised hammer-style pollen press and the powdered sample was added to a pre-weighed vial. In Newcastle University, these samples were reweighed, placed in aluminium vials and burned in a muffle furnace for 2 hours at 450 °C to remove organic matter. After being transferred to a desiccator to cool down to room temperature, the samples were reweighed to determine the weight of non-organic matter. Ash free dry weight was standardised to tissue surface area of each branch based on the length and midpoint width assuming the area of a cylinder, and colony tissue biomass values were assigned as averages across the three branches per colony.

Differences in colony size and tissue biomass (response variables) between thermal refugia and hotspots (2-level fixed effect) were tested using linear mixed effects models under log-link, accounting for site-level differences by fitting a random intercept by site. The effect on heat tolerance (response variable, DHW_50_) of the interaction between either (a) colony size (log-transformed) and thermal regime or (b) tissue biomass (log-transformed) and thermal regime was also tested using linear mixed effects models, accounting for site-level differences using random intercepts. For these interaction models, backward stepwise model selection was used to remove non-important predictors with P values > 0.1, leaving the final models as (a) DHW_50_ ∼ log_tissue_biomass × region + (1|site), and (b) DHW_50_ ∼ log_colony_size + (1|site).

## 3. Results

### 3.1. Persistent thermal refugia and hotspots

Here, we evaluated local-scale thermal stress regimes among the reefs of a small remote archipelago using a 36-year satellite record of sea surface temperatures (SST). The Republic of Palau serves as an ideal case study as it represents more manageable-relevant spatial scales an order of magnitude smaller (<150 km long) than regions previously linked to gradients of thermal exposure and coral heat tolerance (1000s km^4,9,10^). At a first assessment, Palau’s coral reefs appear to have broadly similar historic satellite-based thermal environments, with warm season SST climatologies differing by < 0.1 °C. However, distinct spatial trends emerge in the historic exposure of reefs to accumulated heat stress, measured as degree heating weeks (DHW). DHW captures the intensity and duration of marine heatwaves, where significant bleaching and subsequent mortality are expected beyond DHWs of 4 and 8 °C-weeks, respectively^21^. Marked spatial patterns are evident by ranking reefs in terms of DHW annually (*i*.*e*., 36 maps of spatial DHW rankings, Fig. 1A), and then plotting the relative thermal environment of reefs on a thermal consistency biplot based on two metrics (Fig. 1B). The average DHW rank of a reef across all years (x-axis) captures the typical heat stress it receives relative to others, and the standard deviation of spatial DHW ranks through time (y-axis) captures the temporal consistency of a reef’s relative thermal environment (Fig. 1C). There is a clear gradient of thermal regimes characterised by thermal refugia, hotspots, and thermally moderate reefs (Fig. 1D). Of all the reef grid cells in Palau, 13% emerge as persistent thermal refugia distributed sparsely in the north, an area more exposed to the open ocean, while 18% emerge as persistent hotspots located in a southwestern area associated with high water retention of the shallow inner lagoon (Fig. 1E).

The three strongest marine heatwaves on record in Palau occurred in 1998, 2010, and 2017. During these events, thermal refugia have been consistently exposed to less-intense heat stress than hotspots (2 °C-weeks less on average), despite absolute DHW values still reaching 4–6 °C-weeks, or NOAA Bleaching Alert 1^21^. While local-scale thermal refugia in Palau are not immune to marine heatwaves, they experience less intense heat stress when such events do occur. Cool years without any accumulated heat stress (*i*.*e*., DHW of zero) have been far more frequent at thermal refugia (8-12 years since 1980) than at hotspots (only 3-4 years since 1980), which have been subjected more frequently to years with low-level heat stress. Notably, a small proportion of reefs located at the southern tip of Palau showed the highest interannual variability of heat stress exposure rankings (DHW percentiles) but with a moderate mean DHW rank through time (Fig. 1D, peak of the hump-shaped pattern). In theory, this environmental phenomenon could come about for a reef which behaves as a hotspot in some years and a thermal refuge in others. However, in this case, the highest and lowest DHW percentiles occurred in low heat stress years when DHW was very stable across Palau, suggesting they have moderate thermal regimes.

### 3.2. Hotspots show higher and less variable resistance to mass bleaching

Given historical patterns of mass coral bleaching and marine heatwave stress, we quantified spatial trends in the effect of contrasting thermal regimes on assemblage-wide bleaching severity. As expected^7,22^, DHW was an important predictor of coral bleaching in Palau (Bayesian DHW slope: 0.362 [0.287-0.439], mean and 95% credible intervals). For a given heat stress dosage, the probability of severe assemblage-wide bleaching was highest in the north of Palau. This is concurrent with observations of more severe bleaching in the north in 2010^23^. This result aligns closely to the delineation of thermal refugia and hotspots, with significantly less-severe bleaching predictions at hotspots (Fig. 2D). This could come about through a higher abundance of either heat tolerant coral taxa or heat tolerant individuals for certain species. Moreover, the variability of severe bleaching predictions for a given DHW were higher at thermal refugia than hotspots, with 2.7-fold greater standard deviation, 2.3-fold greater range, and 3.1-fold greater interquartile range of spatial random field (SRF) values (Fig. 2D).

**Figure 2.**
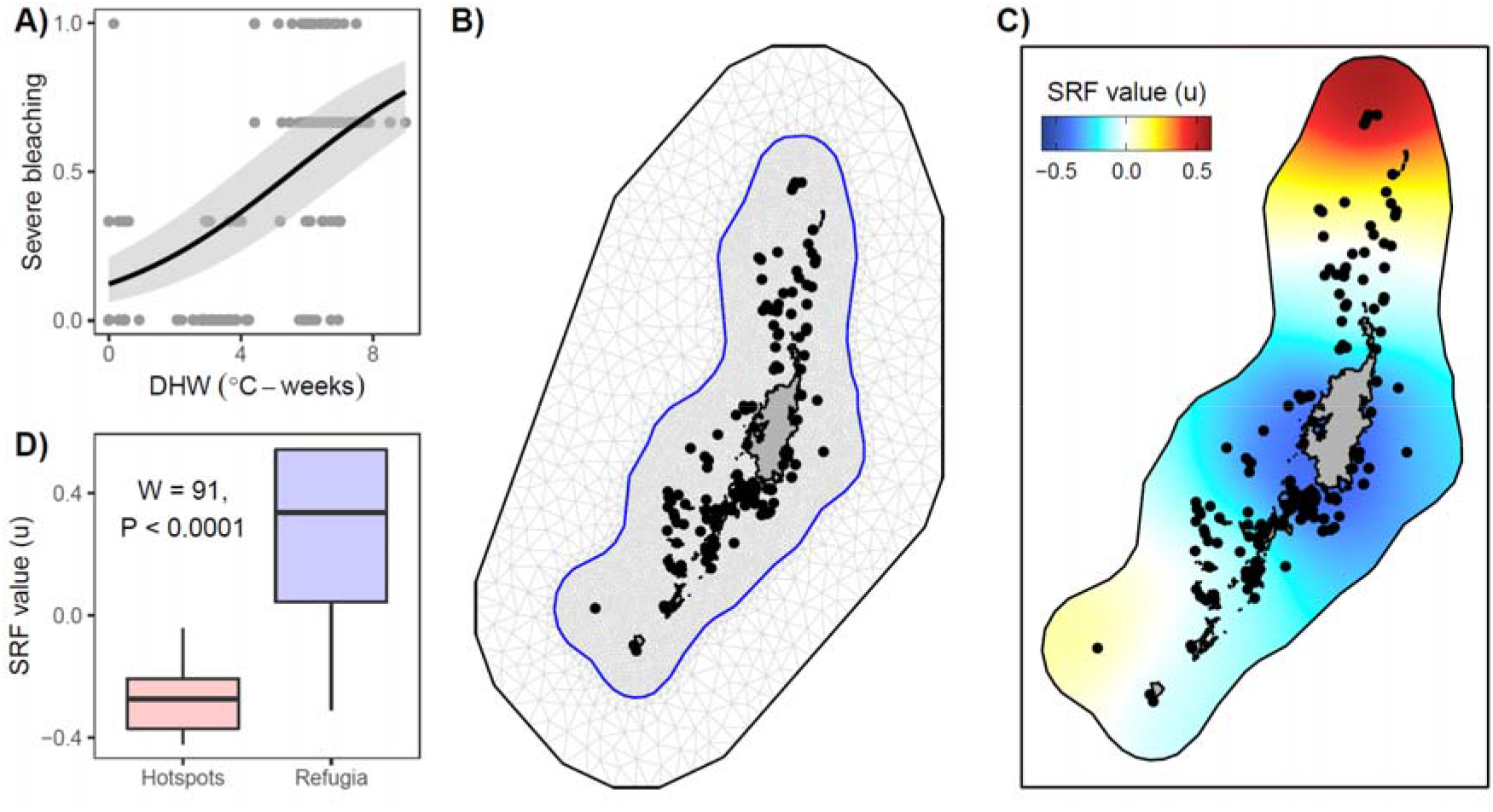
Assemblage-wide bleaching risk across Palau based on historic bleaching survey data and satellite-derived heat stress. (A) The predicted effect of DHW on severe coral bleaching is shown as the mean and 95% credible intervals with notable overprediction and underprediction for some bleaching observations. (B) Spatial correlated uncertainty in bleaching predictions is calculated across a high-resolution Delaunay triangulation mesh of the study area comprising 6,210 nodes. (C) The spatial random field (SRF) reports the spatial correlated uncertainty as u values (see linear combination equation), showing higher bleaching susceptibility for a given DHW dosage in red (underpredictions) and higher bleaching resistance in blue (overpredictions). (D) Thermal refugia have higher mass bleaching susceptibility for a given heat stress dosage (SRF values) than hotspots (Wilcoxon sum rank test) and a higher diversity of responses.

### 3.3. Thermal refugia show higher and more diverse species-specific heat tolerance

Because differences in species composition between reefs could mask any influence of selection or acclimatisation on assemblage-wide heatwave responses, we restrict our spatial comparison of heat tolerance to a single species, *Acropora digitifera*, a common Indo-Pacific reef building coral^17,24^. We determined individual heat tolerance by experimentally subjecting coral fragments from hotspots and thermal refugia to a long-term 5-week marine heatwave experiment, reaching a final DHW of 16.2 ± 0.2 °C-weeks (mean ± SD across tanks). We found marked differences in *Acropora digitifera* heat tolerance between thermal regimes.

The progression of bleaching and mortality responses among colonies was highly variable. Heat tolerance, measured as DHW_50_ (the DHW dosage causing a 50% bleaching and mortality response or BMI=0.5), was significantly higher at thermal refugia (Fig. 3B). Compared to colonies from hotspots, those from thermal refugia could withstand an additional 0.7 °C-weeks of heat stress before the onset of bleaching and mortality (Fig. 3A). This contrasts with our finding of lower assemblage-wide heat stress tolerance at thermal refugia.

**Figure 3.**
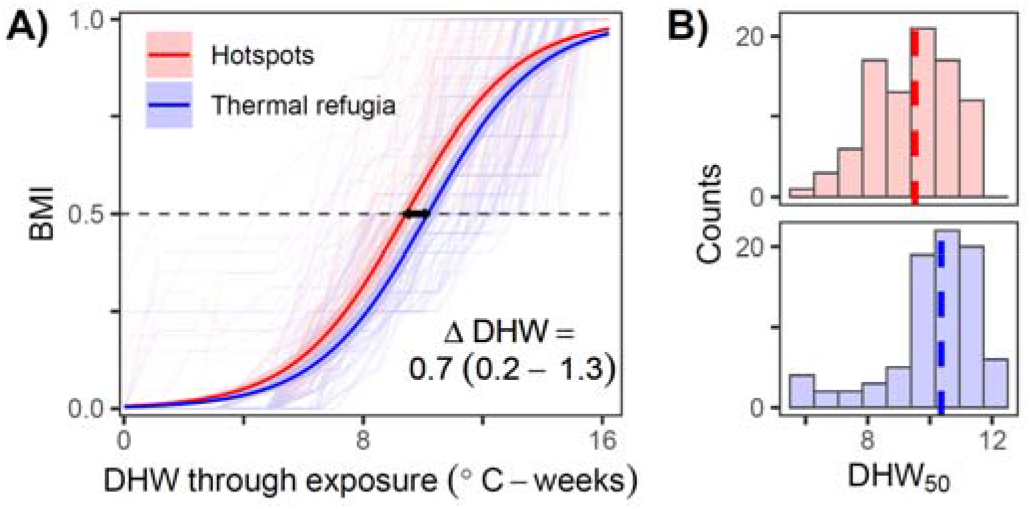
Differences in *Acropora digitifera* heat tolerance between hotspots (red) and thermal refugia (blue). (A) Bleaching and mortality index (BMI) trajectories are shown for each individual coral (faint lines) throughout a 5-week marine heatwave experiment relative to degree heating weeks (DHW), with a significant difference between thermal regime intercepts (mean ± 95% confidence intervals, n=174 colonies) (GLMM, *P* < 0.001). (B) Distribution of individual heat tolerances (DHW_50_) are shown for each thermal regime with a significantly higher mean heat tolerance (dashed lines) at thermal refugia (LM, *P* = 0.009).

Heat tolerance variability was greater within than between thermal regimes. For hotspots, there was a ΔDHW of 4.1 °C-weeks (2.5 – 5.8 °C-weeks 95% CI) between the bleaching and mortality responses of the least- and most-tolerant 10% of colonies (Fig. 4A). This translates to a categorical shift in the NOAA Bleaching Alert system (*e*.*g*., Alert Level 1 to 2). However, heat tolerance variability was 25% greater at thermal refugia (Fig. 4E), with a ΔDHW of 5.2 °C-weeks (3.5 – 7.2 °C-weeks 95% CI) between the upper and lower deciles of the population. There was an indication that higher variability at thermal refugia was due to the presence of both the most-sensitive and most-tolerant colonies, yet we detected no significant differences between thermal regimes for the least-tolerant (GLMM, *P* = 0.79) or most-tolerant decile subsets (GLMM, *P* = 0.19).

**Figure 4.**
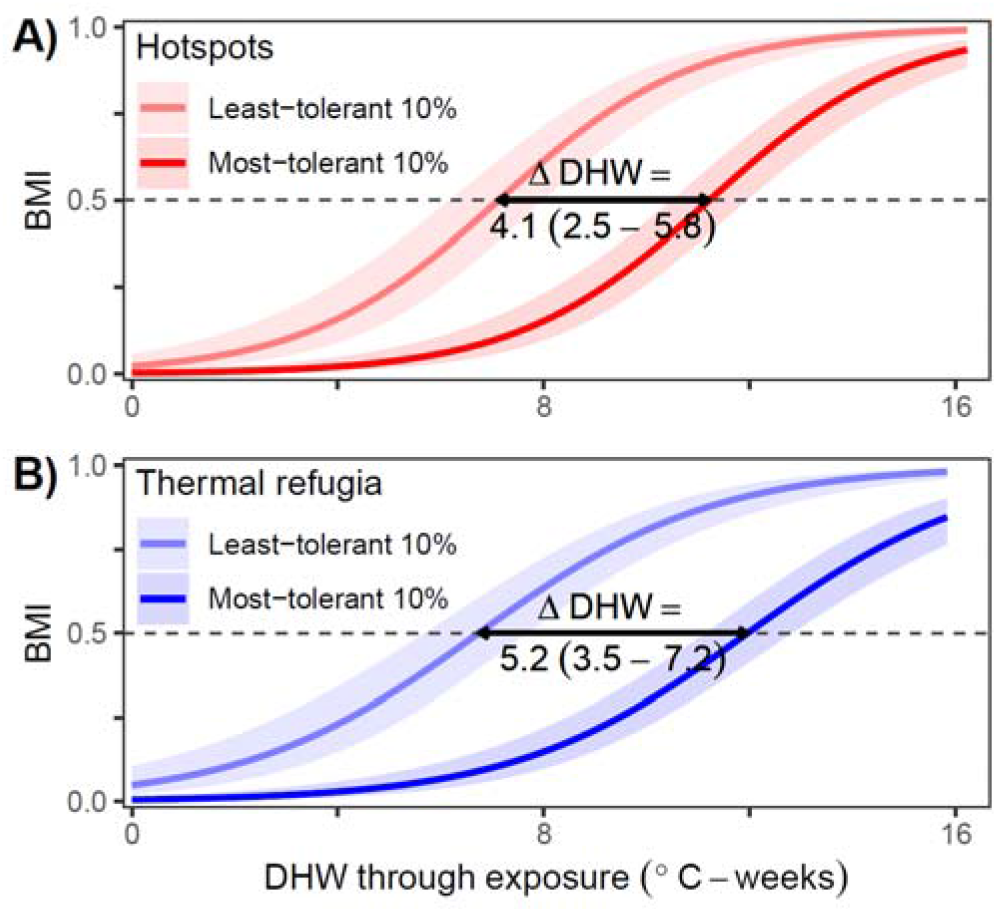
Within-population variability in heat tolerance for *Acropora digitifera* corals at hotspots (A) and those at thermal refugia (B), expressed as differences between the least-tolerant decile (paler shading) and most-tolerant decile (darker shading) of the population (n=9 colonies for each subsample in each thermal regime) in terms of DHW tolerance (ΔDHW, mean and upper and lower 95% confidence limits) evaluated at BMI = 0.5 (grey dashed line).

### 3.4. Phenotypic drivers of species-specific heat tolerance

As an indication of potential phenotypic drivers of heat tolerance, we also assessed colony size and tissue biomass. Sampled *A. digitifera* colonies had significantly larger sizes at thermal refugia than at hotspots (Fig. 5A). A positive correlation was found between colony size and heat tolerance (Fig. 5C), albeit with a high level of uncertainty (*P* = 0.096), such that larger colonies had a higher heat tolerance. For tissue biomass, while no significant difference between thermal regimes was detected (Fig. 5B, *P* = 0.84), a positive correlation with heat tolerance was found at thermal refugia (Fig. 5 D, *P* = 0.015). However, this trend was absent at hotspots (interaction term, *P* = 0.037).

**Figure 5.**
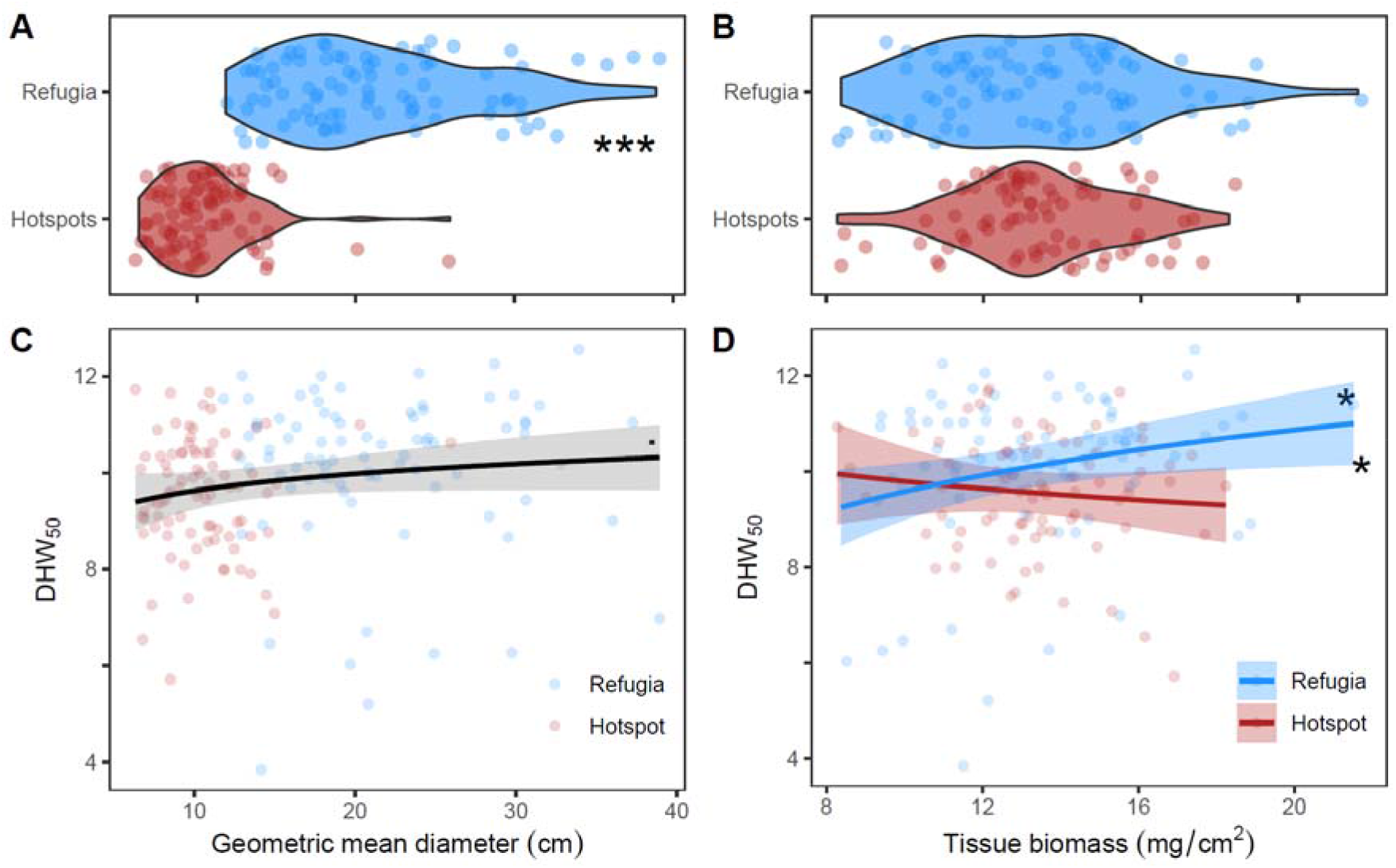
Phenotypic drivers of heat tolerance between thermal regimes. (A,B) Differences in trait values between thermal refugia (blue) and hotspots (red), and influence of traits on heat tolerance (C,D). Each test was performed using separate linear mixed effect models (LMMs), accounting for site-level differences by fitting a random intercept for each site. LMMs with hypothesised interactions were reduced using stepwise backward selection. (A) Coral colonies were significantly larger at thermal refugia (P < 0.001). (C) There was a positive association between colony size and heat tolerance associated with a moderate level of uncertainty (P = 0.096), and with no interaction between thermal regimes (P > 0.05). (B) No differences in mean tissue biomass were detected between thermal regimes (P = 0.84). (D) Tissue biomass was positively associated with colony heat tolerance at thermal refugia (refugia slope, P = 0.015), but this effect was not present at hotspots (interaction, P = 0.037) where the trend did not differ significantly from the null expectation of a slope equal to zero (P > 0.05). P value significance is given by black symbols for P < 0.1 & > 0.05 (.), P < 0.05 (*), P < 0.01 (**), P < 0.001 (***).

## 4. Discussion

Theory and evidence show that large geographic-scale gradients in thermal exposure play a major role in determining the heat tolerance of marine and terrestrial organisms, through mechanisms including acclimatisation and selection. However, it remains unknown whether this theory is robust over the smaller spatial scales at which organisms and effective management operate. Here, we show that this theory does not necessarily hold true when applied to coral heat tolerance. We identified persistent local-scale thermal refugia and hotspot reefs within 60km of one another at a remote Pacific archipelago. Given their exposure to heat stress conditions, hotspots are expected to be associated with higher coral heat tolerance. At the level of the coral assemblage, patterns in historic mass bleaching severity support these expectations, in line with the fact that refuge coral assemblages are naive to marine heatwave stress. However, when comparing the heat tolerance of *Acropora digitifera* between thermal refugia and hotspots, we found the opposite trend, with higher heat tolerance at thermal refugia. Furthermore, we found a greater diversity of bleaching responses at thermal refugia for both the assemblage and species levels. Ultimately, our study challenges the theory that, compared to thermal refugia, hotspot reefs should harbour higher coral heat tolerance. We also show that spatial patterns in mass coral bleaching do not necessarily predict species responses. Our results highlight the importance of understanding local-scale phenomena to provide actionable information to address global-scale adaptive management problems.

Theory on coral heat tolerance and local adaptation have been developed and tested across large spatial thermal gradients^15^, between reef habitat with extremely different temperature regimes (*e*.*g*., outer reefs versus lagoons^25,26^), and in natural field experiments such as neighbouring backreef tidal pools (<500 m apart)^27^. These studies show that higher temperatures and thermal variability can lead to acclimatisation, selective pressure, and ultimately higher heat tolerance^25,26^. However, at spatial scales that can be mapped by satellite-borne thermal sensors (>1-10km), how coral heat tolerance is influenced by habitat-specific thermal regimes on (*e*.*g*., at outer reefs only) remains poorly understood, despite the importance of microhabitat variability on adaptative capacity more broadly^28^. Our study found contrasting patterns at different levels of biological organisation, with hotspots associated with higher assemblage-wide tolerance but lower species-specific heat tolerance. The assemblage-level trend could be due to local differences in coral species composition (*i*.*e*., reefs with more abundant heat-sensitive taxa like *Acropora* are expected to suffer more severe bleaching impacts^29^). However, this may not be the case for Palau, given widespread temporal stability of *Acropora*^30^. Therefore, these results suggest that spatially varying acclimatisation or adaptation may be occurring for some members of the assemblage. Yet, this has not taken place for the study species, indicating that non-thermal environmental drivers may also influence heat tolerance.

Variability in *A. digitifera* heat tolerance was 25% higher at thermal refugia than hotspots (ΔDHW of 5.2 and 4.1 °C-weeks, respectively). Previous work on *A. digitifera* from a thermally moderate eastern outer reef in Palau detected a ΔDHW of 4.8 °C-weeks^17^ between upper and lower deciles of heat tolerance. Combining this with the results presented here, heat tolerance variability may be negatively correlated to historic heat stress exposure in our study system. The most-sensitive corals were found at thermal refugia but were absent at hotspots, suggesting that thermal selective pressures may be more prevalent at hotspot reefs. Yet the most-tolerant corals were also absent at hotspots, suggesting a potential role of non-thermal selective pressures influencing heat tolerance (*e*.*g*., lower water quality^18,19^). Notably, we also found that heat tolerance variability was widespread. Provided coral heat tolerance has a genetic basis and is heritable^31–34^, this indicates a high potential for coral adaptation to climate change.

Coral heat stress responses are complex and influenced by biological and environmental factors^7,35,36^. Our results show that larger colonies and those with thicker tissue, a proxy of energy reserves, could withstand an additional 1 °C-week of heat stress before bleaching and dying. Larger colony sizes at thermal refugia than at hotspots suggest either a difference in age structure or growth rates. Notably, the removal of colony fragments was easier at thermal refugia, indicating lower skeletal density which has been suggested as a potential trade-off with *A. digitifera* heat tolerance in Palau^37^. Differential heat tolerance between thermal regimes can also be influenced by symbiotic microalgal communities. Certain symbiont taxa (*e*.*g*.,, *Durusdinium* spp.) provide corals with higher heat tolerance^38^.

However, previous studies of Palau’s outer reefs have found *A. digitifera* is ubiquitously dominated by *Cladocopium C40* symbionts with no link to heat tolerance^17^, and four other coral genera also host predominantly *Cladocopium* spp. symbionts^25^. Although we cannot rule out microalgal symbionts as drivers of heat tolerance, these studies suggest that *Durusdinium* spp. is rare on Palauan outer reefs despite prevalence in lagoons^39^, reflecting findings from other Pacific islands^40^. Thus, other mechanisms may provide more parsimonious explanations of heat tolerance, including host genetics or phenotypic plasticity driven by environmental parameters other than temperature.

Despite the sampled areas of reef having the same depth range and aspect, numerous environmental parameters are known to influence Palau’s coral reefs, including wave exposure^41^ and water quality^42^. Such factors can influence coral bleaching^18^ and can be correlated to thermal environments. The unexpected high heat tolerance at thermal refugia, together with other phenotypic and morphological differences between thermal stress regimes, suggest that heat tolerance is not only a function of the thermal environment.

Indeed, the altered colony morphology observed here between thermal regimes suggests that differences in wave exposure, currents or nutritional opportunities exist, which could influence coral heat tolerance. It could prove insightful to categorise physical reef environments – as for the Caribbean^43^ – and then revisit the role of thermal environments on coral heat tolerance. We may yet discover that the theory of higher tolerance existing in warmer and/or more variable regions holds true in physically comparable environments.

Thermal refugia have previously been identified across global^44^ and regional^9^ scales (>100-1000 km). Yet here we find persistent thermal refugia within 60 km of hotspots. Local variability in thermal regimes at the same spatial scales at which corals operate (see ^45,46^) will likely be important for coral reef persistence and the feasibility of novel reef management approaches. For instance, climate-smart marine protected area networks that aim to boost adaptation rates^15,47,48^ could benefit from local-scale variability in thermal regimes detectable from satellites, by protecting key dispersal links between reefs. Similar benefits could apply to assisted evolution, an interventionist approach to boosting natural rates of climate adaptation by propagating more tolerant individuals. For instance, translocating coral colonies from reefs with different characteristics may be more feasible if donor reefs are closer to recipient reefs (e.g., lower costs, less risks of contamination, fewer permitting issues). However, the science underpinning these novel management approaches is still in its infancy, and they have yet to be implemented in practice.

A dilemma for designing assisted evolution interventions emerges in the choice between assisted geneflow (based on translocating corals from reefs expected to harbour heat tolerant genotypes^49^) versus selective breeding (based on identifying heat tolerant parental genotypes on any reef and out-planting their offspring locally^24^). For assisted geneflow, the main investment would be in translocation costs, whereas for selective breeding, higher investment would be required for identifying heat tolerant genotypes on any given reef. Since coral heat tolerance was spatially widespread in Palau (this study and ^50^), there may be a higher return-on-investment for selective breeding over assisted geneflow. However, this may only be applicable in the context of remote coral reef systems with limited habitat-specific environmental gradients, and populations with high genetic diversity (*e*.*g*., many remote Pacific reefs). The relative feasibility of selective breeding and assisted geneflow may be different for larger reef systems that span considerable thermal gradients (see ^4,9,10^), or for populations that have already suffered substantial declines with low remaining genetic diversity (*e*.*g*., *Acropora palmata* in the Caribbean^49^).

Locations with consistent exposure to specific climatic stressors are thought to harbour individual organisms with higher tolerance to such stressors through acclimatisation and selection. While this theory holds true across broad climatic gradients^4,5^, we find the theory can break down at the intrapopulation scale. We demonstrate this for a remote coral reef system and individual coral tolerance to marine heatwave stress and bleaching. In the context of small coral reef systems with limited gradients in thermal exposure, our study challenges the paradigm that corals at thermal refugia are necessarily maladapted to high temperature conditions^15^. Moreover, by focussing on intensive intrapopulation sampling at local spatial scales we have shown that there is considerable innate potential for adaptation to climate change. Our findings highlight the importance of scale when understanding heterogeneity in biological responses to climate change, and have implications for other ecological systems (*e*.*g*., forests and seagrass meadows). Knowing that large-scale theory is not necessarily robust at small intrapopulation scales, it follows that a better understanding of local-scale ecological phenomena is needed to aid in the design of climate-smart initiatives capable of addressing global-scale adaptive management problems.

## Data and code availability

All original data and code (R version 4.0.2, GNU Bash version 5.0.16(1) and CDO version 1.9.9rc1) have been deposited at 10.25405/data.ncl.22731194 (available before publication at https://figshare.com/s/faac387cb999778055cc to be completed). All datasets analysed are publicly available as of the date of publication. Any additional information required to reanalyse the data reported in this paper is available from the lead contact upon request.

## Acknowledgements

This works was supported by a UKRI Mitacs Globalink grant to L.L., J.R.G, and S.D. (NE/T014547/1), an International Coral Reef Society Ruth Gates Fellowship to L.L., an IDEAWILD fieldwork equipment grant to L.L. (LACHPALA1219), the Natural Environment Research Council’s ONE Planet Doctoral Training Partnership Studentship (NE/S007512/1) to L.L., and a European Research Council Horizon 2020 project CORALASSIST (725848) to J.R.G. We also thank Dr. Harmony Martell, Dr. Pedro Gonzalez-Espinosa, and Xinru Li for their thoughts on this work, PICRC staff for supporting our research, Arius Merep for help with aquarium tank construction, and the boat operators Geory Mereb, and Nelson Masang.

## Author contributions

L.L., S.D.D., A.H., P.J.M., and J.R.G. conceived and designed the study; L.L. developed the computer code; L.L., S.D.D., P.J.M., J.B., and J.R.G analysed the results; L.L., E.B., L.B., D.B., J.B., R.T.C, A.J.E., A.H., H.M.M, E.S., A.W, and J.R.G completed the marine heatwave experiment and fieldwork; L.L. wrote the first draft of the paper; All authors critically revised the manuscript.

