## Supplementary Materials for "Unexpectedly high coral heat tolerance at thermal refugia"

**Supplementary Text 1.** A small proportion of reefs located at the southern tip of Palau showed the highest interannual variability of heat stress exposure rankings (DHW percentiles) but with a moderate mean DHW rank through time (Fig. 1D, peak of the hump-shaped pattern). In theory, this environmental phenomenon could come about for a reef which behaves as a hotspot in some years and a thermal refuge in others. However, in this case, the highest and lowest DHW percentiles occurred in low heat stress years when DHW was very stable across Palau, suggesting they have moderate thermal regimes.

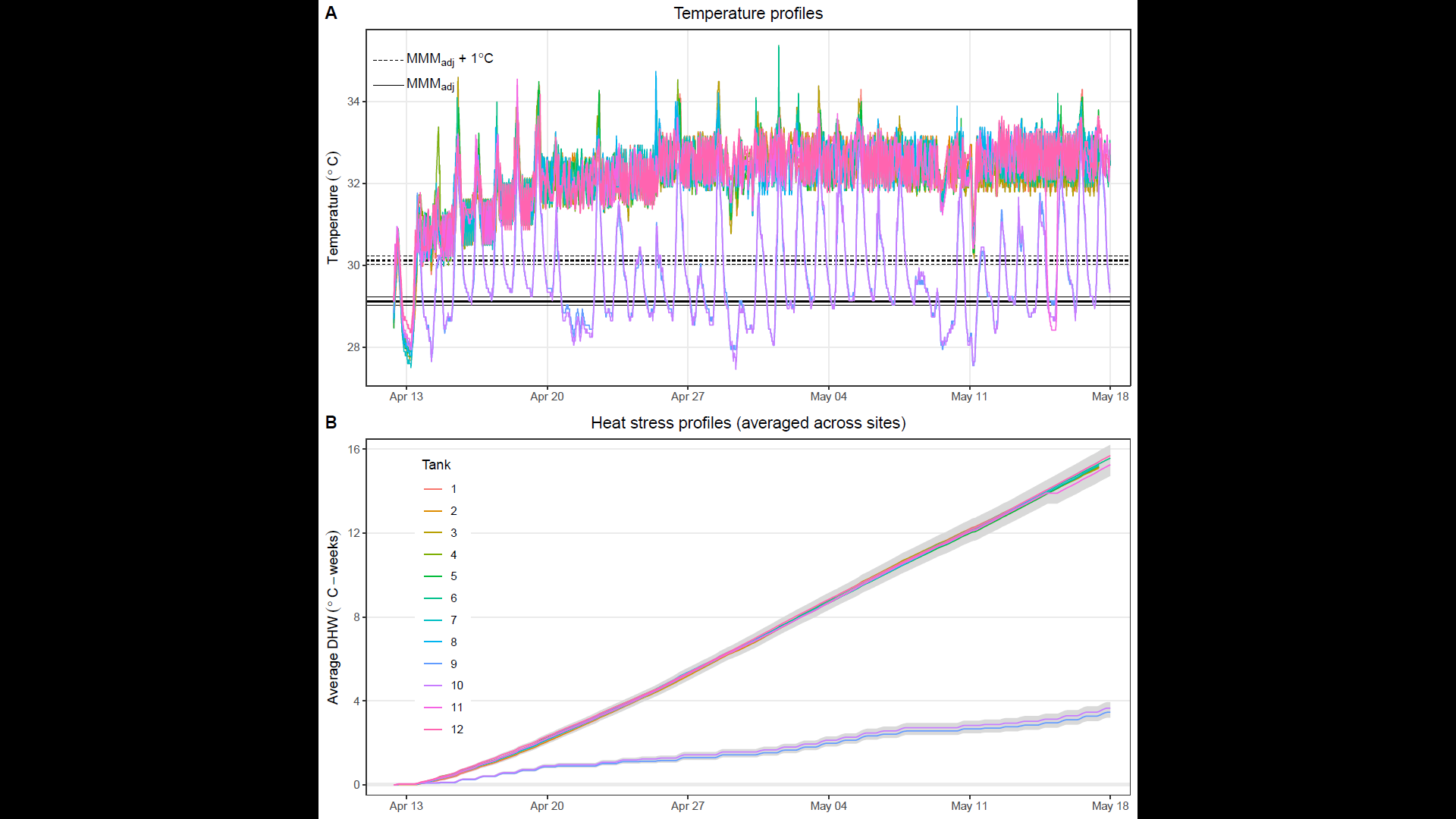

**Figure S1. Simulated marine heatwave experiment.** (A) Temperature profiles for the ten replicate heat stress tanks (Tanks 1–8, 11, and 12) and procedural control tanks (Tanks 9 and 10), sharing the colour legend with panel B. The local climatological baseline (MMM_adj_) and the corresponding stress accumulation threshold (MMM_adj_ + 1 °C) are shown for each of the 6 coral collection sites. Specific values for each site are shown in Table S1. (B) The average accumulated heat stress profile (DHW – degree heating weeks) is shown as the average across all sites in each experimental tank (colour legend). The shaded region shows the total range of absolute DHW across all sites and tanks.

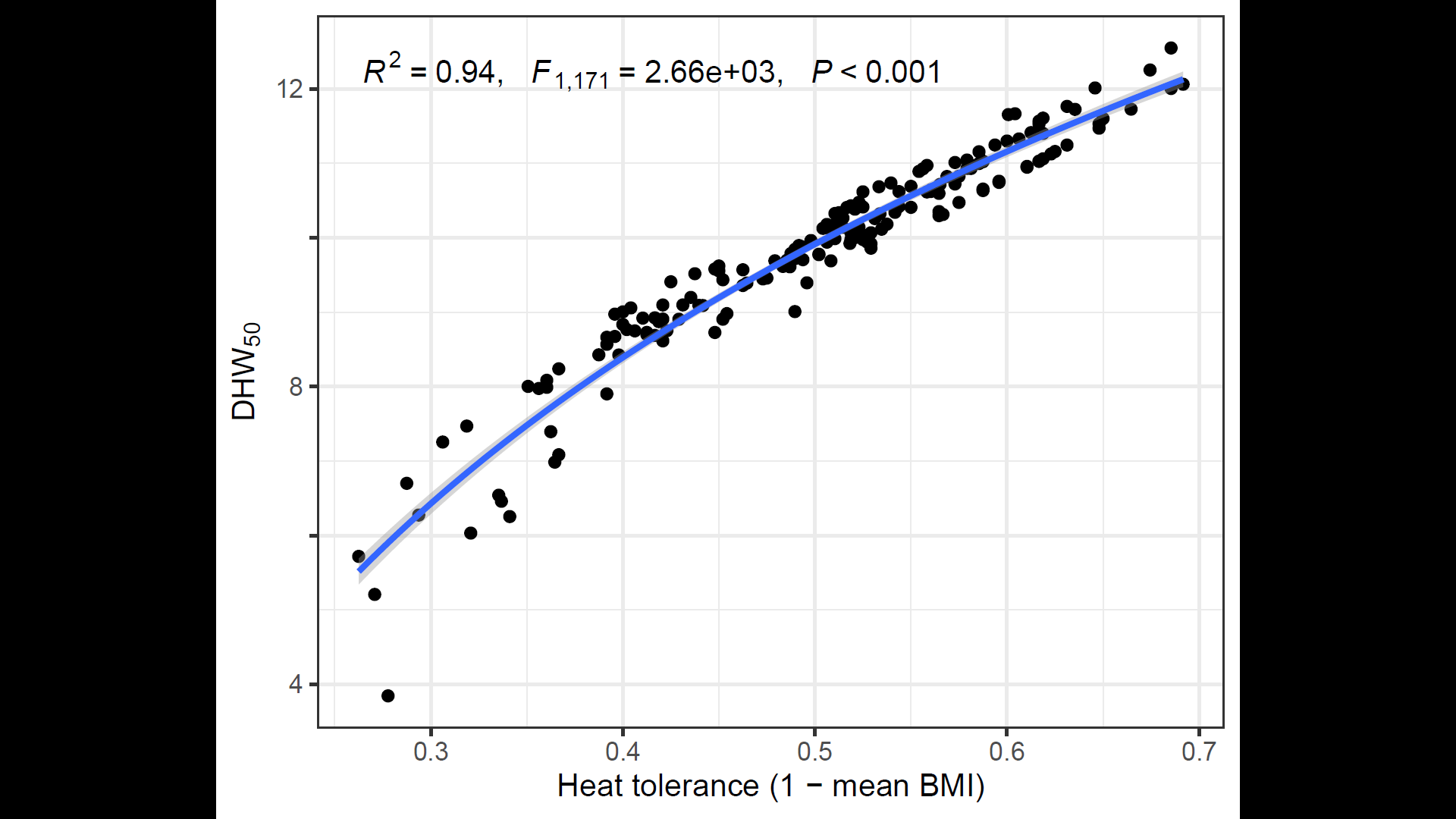

**Figure S2.** Relationship between inverse mean BMI metric of heat tolerance and DHW_50_, the heat stress dosage at which a colonies response is predicted to pass a BMI of 0.5.

**Table S1.** Simulated marine heatwave experiment. Associated metadata following ^66^.

| **Metadata** | **Conditions or methods** |
| --- | --- |
| Coral collection | **Latitude and longitude:**  For 3 hotspot sites (H) and 3 refugia sites (R) in decimal degrees  R1: 7.965, 134.625; R2: 7.989, 134.659; R3: 7.826, 134.630  H1: 7.374, 134.279; H2: 7.410, 134.330; H3: 7.290, 134.241 |
|  | **Collection depth:** between 1 and 5 m depth |
|  | **Collection dates:** April 3^rd^ – April 5^th^ 2022 |
|  | **Coral species:** *Acropora digitifera* |
|  | **Coral morphology:** corymbose |
|  | **Symbiodinaceae for all colonies:** NA |
|  | **Acclimation post collection prior to experiment:** 7 – 10 days |
| Experimental design | **Name of location:** Palau International Coral Reef Center, Palau |
|  | **Bleaching stress temperature period:** April 13^th^ – May 18^th^ 2022 |
|  | **System type:** outdoor flow through system |
|  | **No. tanks per treatment:** 10 heat stress & 2 procedural control tanks |
|  | **No. coral genets (colonies) per treatment:** 178 with 6 fragments/colony |
|  | **No. coral genets (colonies) per tank within treatments:** 75 – 97 |
| Experimental temperature conditions | **Heat stress temperature above MMM per treatment:** approximately + 3.5 °C above the local climatological baseline, MMM_adj_ (see baseline temperature description below) |
|  | **Control temperature:** 29.79 ± 1.18 °C |
|  | **Baseline temperature:** CoralTemp MMM was 29.2 °C for all 6 sites but was adjusted based on the relationship between satellite sea surface temperatures and *in situ* temperatures from each site, resulting in MMM_adj_ values of 29.0, 29.2, and 29.1 °C for H1, R3, and the remaining 4 sites, respectively. The stress baseline for accumulating DHW was set to MMM_adj_ + 1 °C |
|  | **Temperature ramp-up rate:** Compared to the MMM_adj_ (among site average of 29.1 °C), temperatures in heated tanks were adjusted by + 1.4 ºC on day 1 to a temperature of ~30.5 ºC; by + 0.5 ºC on days 3, 5, 7, and 13 to a temperature of ~32.5 ºC |
|  | **Duration at heat stress temperature:** 35 days |
|  | **Temperature modulation:** diurnal |
| Other experimental conditions | **Light conditions:** 400 μmol photons m^−2^ s^−1^ |
|  | **Light cycle:** 12 h:12 h diurnal cycle |
|  | **Water flow velocity:** 7.2 L/hour |
|  | **Tank turnover:** 2 times per day |
|  | **Sea water filtration:** 50µm filtered |
|  | **Sea water source:** natural directly from the reef |
|  | **Salinity:** 35‰ |
|  | **Feeding:** natural feeding from plankton in the filtered seawater |
